# *Streptococcus pneumoniae* fratricide-induced cell lysis does not require the type IV competence pilus

**DOI:** 10.1101/2025.11.06.685197

**Authors:** Anna K. Borowska, David T. Eddington, Donald A. Morrison

**Affiliations:** Department of Biomedical Engineering, University of Illinois Chicago, Chicago, IL, 60607, USA; Department of Biological Sciences, University of Illinois Chicago, Chicago, IL, 60607, USA

## Abstract

*Streptococcus pneumoniae*, a bacterium that becomes competent naturally, can undergo transformation and fratricide during which different genes, including antibiotic-resistant ones, can be transferred. A Type IV pilus (T4P) is involved in the DNA take up during transformation or fratricide-induced (fratricide = same species cell killing) gene transfer. Here, contrary to our expectations, we show that competence T4P is not required in fratricide-induced cell lysis. We have utilized pilus⁺ and pilus⁻ attackers, and a β-galactosidase^+^ (β-gal^+^) victim for our experiment. Transformation, gene transfer efficiency, and cell lysis measured by release of β-gal^+^ were tested at different optical cell densities (OD) under competence-induced (+CSP) or non-competence (-CSP) conditions. We have optimized experiment conditions to observe cell lysis, which only occurred in the media that do not contain choline (inhibits cell lysis). As expected, we have observed transformation and gene transfer with the pilus^+^ strain, but none of those two processes with the pilus^-^ strain. In the case of β-gal release during fratricide, we have observed β-gal activity in both cases whenever the pilus^+^ or pilus^-^ strain was mixed with the β-gal^+^ victim. Our data indicate that type IV competence pilus, while crucial for DNA take up during transformation fratricide, is not needed during cell lysis.

## INTRODUCTION

*Streptococcus pneumoniae*, commonly known as pneumococcus, is a human pathogen responsible for infections such as pneumonia, bacteremia, and meningitis (1–6). According to WHO data from 2019, it is linked to approximately 300,000 deaths among children under five years of age worldwide (7). Antibiotic usage is a primary way to treat pneumococcal infections. However, developing additional protection systems like vaccines is essential (8), due to pneumococcal diseases spreading, and existing treatments are proving insufficient. This spread is caused by horizontal gene transfer (HGT), particularly through competence - a metabolic state influenced by cell density - and transformation (9,10). During these processes, *S. pneumoniae* cells become competent, killing neighboring cells and incorporating their DNA (11).

As mentioned above, *S. pneumoniae* cells can become competent and transform. Griffith first described transformation in 1928 (12) and later in 1944 Avery demonstrated that genetic material released from capsule-containing virulent cells was taken up by an avirulent one (13). In *S. pneumoniae,* such transformation connects to a specific metabolic state, known as competence, a natural condition that can be activated only under certain circumstances, and has been described by many (14–19).

Pneumococcal competence and transformation depend on a quorum sensing mechanism (20), controlled by two types of genes: early and late *com* genes. Briefly, the early *com* genes are *comA–comE* and *comX,* and one of their main products, specifically from *comC*, is a competence signaling peptide (CSP), which is responsible for initiating the competence state. The products of early *com* genes activate late *com* genes, whose expression results in the expression of a competence type IV pilus (T4P) (genes *comGA–comGG*) and other machinery, including EndA, ComEA, ComFC, SsbB, and DprA. This machinery facilitates the uptake of single-stranded DNA into the cell and its integration into the bacterial genome (21).

The late *com* genes *comGA – ComGG,* encode T4P in *S. pneumoniae* (22). These gene products have distinct roles in pilus formation: *ComGA* encodes a putative ATPase involved in pilus assembly, *ComGB* encodes a polytopic membrane protein, *comGC* encodes the main pilin subunit and *comGD-comGG* encodes prepilins, with function remained unclear for a long time (22). Recently, a group of Pelicic (23) showed in Streptococcus sanguinis that electropositive residues of two minor pilins, ComGD and ComGF, are involved in DNA binding during transformation. Those proteins are conserved among bacteria with one membrane, indicating that ComGD and ComGF are involved in DNA binding in *S. pnuemoniae* (23).

The T4P in *S. pneumoniae* was first visualized in 2013 by Laurenceau et al. (22). Later, data showed that the T4P of *S. pneumoniae* can extend and bind DNA. *S. pneumoniae* T4P is not the first DNA-binding pilus described in gram-positive bacteria. Previously, *Bacillus subtilis* was found to possess a pseudopilus, that comes from the same operon *comGA – ComGG* as *S. pnuemoniae* (24). Scientists thought that *B. subtilis* pseudopilus could not extend to the environment but could bind DNA and deliver it to the space between the cell wall and cell membrane (25). Recently, studies showed that *B. subtilis* also has T4P, extending, retracting, and binding DNA (26–29).

A distinct process involving a pilus in cell contact is called conjugation, observed in *E. coli* and carried out by an F factor. The F Pilus serves two functions in this process: bringing pair of cells together and participating in DNA transport between the mating cells (30,31). Previous data show that the F pilus sporadically extends and retracts, bringing pair of cells together during retraction (32). In contrast, in 2023, Goldlust et al. (30) showed that the conjugation F pilus serves as a channel for DNA during conjugation between cells that are not in direct physical contact (30). Research indicates that the F-pilus promotes harmful bacteria like *E. coli* (in a tight and distant conjugation). Still, there is no knowledge about pili involved in conjugating gram-positive bacteria (33,34).

In 2002, a group from Norway reported a new phenomenon occurring between *S. pneumoniae* cells, which they named fratricide (35). In this process lysis of the target cells (competence-uninducible ones) was observed, after mixing competence-inducible cells with competence-uninducible ones of the same species. They showed β-galactosidase (β-gal) enzyme release (cell lysis) and DNA uptake from lysed cells by the competent cells. In addition to that they concluded that cell lysis, is influenced by cell phase growth, as it occurs during the logarithmic growth phase (OD = 0.2 – 0.55) (35), since once the cells enter the stationary phase, they become unresponsive to CSP (36,37).

The same group highlighted the importance of LytA in cell lysis (35). Over the years, it has become evident that fratricide-induced cell lysis requires three key proteins: LytA, LytC, and CbpD. The central player in this process is CbpD (Choline-binding Protein D), which functions as a murein hydrolase and plays a significant role in cell lysis (37,38). The key difference between LytA, LytC, and CbpD is their expression patterns. Expression of LytA and LytC is constitutive, while CbpD is expressed exclusively in competent cells, as CbpD is a late competence gene. Studies show that CbpD can activate LytA and LytC to facilitate cell lysis. Earlier reports indicate that CbpD remains bound to the cell wall during lysis, suggesting that direct cell contact is required for fratricide, and potentially this direct contact can involve the recently described T4P (22,34).

Since it has been suggested that CbpD may require direct cell–cell contact to induce cell lysis—and given the lack of reports implicating the T4P in fratricide-associated lysis or in conjugation, both of which involve close cell contact—we propose that the T4P may play a role in initiating this contact in *S. pneumoniae*. As previously described, the T4P is known to be involved in DNA uptake during the transformation process. However, it remains unclear whether this structure also supports the activity of other proteins involved in fratricide-induced cell lysis.

## RESULTS

### 1. Strain construction

To investigate the potential involvement of the T4P in cell lysis, we first prepared the bacterial strains required for the assay. We needed three distinct strains: (I) a strain expressing β-galactosidase (β-gal) that is unable to develop competence, referred to as the target or victim cells; (II) a strain capable of becoming competent and producing pili (designated pilus⁺); and (III) a strain that can become competent but does not produce pili (designated pilus⁻).

To generate non-competent target cells, we introduced the β-gal gene into a competent strain and subsequently rendered it non-competent. The competent parental strain was essential, as only competent cells can efficiently take up exogenous DNA from the environment.

β-gal was selected as the reporter enzyme due to its cytoplasmic localization— attributable to the absence of a signal peptide, which prevents its secretion outside the cell (39) and many reliable, well-characterized detection methods. Although the chromogenic substrate o-nitrophenyl-β-D-galactopyranoside (ONPG), is commonly used for β-gal detection, we opted for a fluorescent-based assay to improve sensitivity. Specifically, we employed fluorescein di-β-D-galactopyranoside (FDG), which yields a fluorescent signal upon enzymatic cleavage by β-gal, producing fluorescein. Higher sensitivity of FDG led to its use compared to chromogenic substrate (40).

We inserted the β-gal gene into the *hirL* locus (Figure 1), which has high expression during exponential growth and disruption does not adversely affect cell growth or competence (35,37). Figures 3a and 3b represent the results of insert generation, introduced into CP2000 competent cells to replace *hirL* with the *lacZ* gene. We designated new obtained strain CP2600 (β -gal^+^).

**Figure 1.**
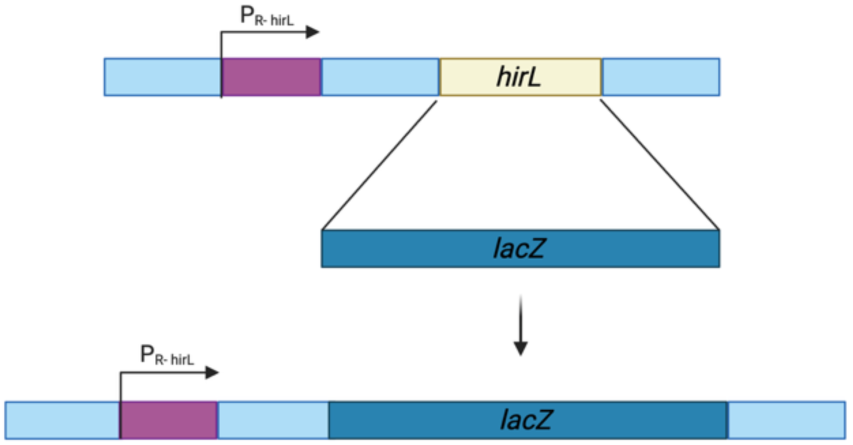
Schematic representation of insertion of *lacZ* gene into *hirL* position. LEGEND: P_R-hirL_ - promotor of *hirL* gene, *hirL* - *hirL* gene, *lacZ* - LacZ gene, encoding β-gal.

After we were sure that our clone was β-gal^+^, we needed to remove competence. Our group previously utilized a non-competent strain called CP2215. It carries a spectinomycin-resistance gene inserted into the *comE* gene, cause its cells to be unable to respond to CSP, since it plays a crucial role in competence cascade activation (Figure 2). We designated our “victim” or “target” strain CP2601 (β-gal^+^, Spc^R^, Figure 4).

**Figure 2.**
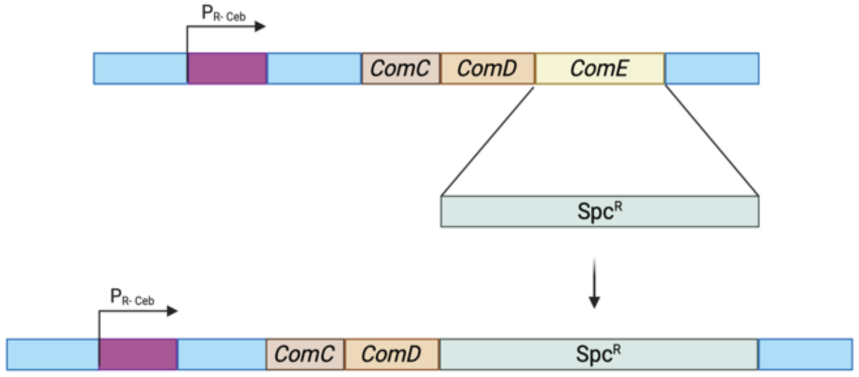
Schematic representation of insertion of *spc* gene into *comE* gene position. LEGEND: P_R-Ceb_ - promotor of ComCDE, ComC – ComD – Comc & ComD genes; *ComE* – *comE* gene, *Spc^R^* – Spectinomycin-resistance gene.

**Figure 3.**
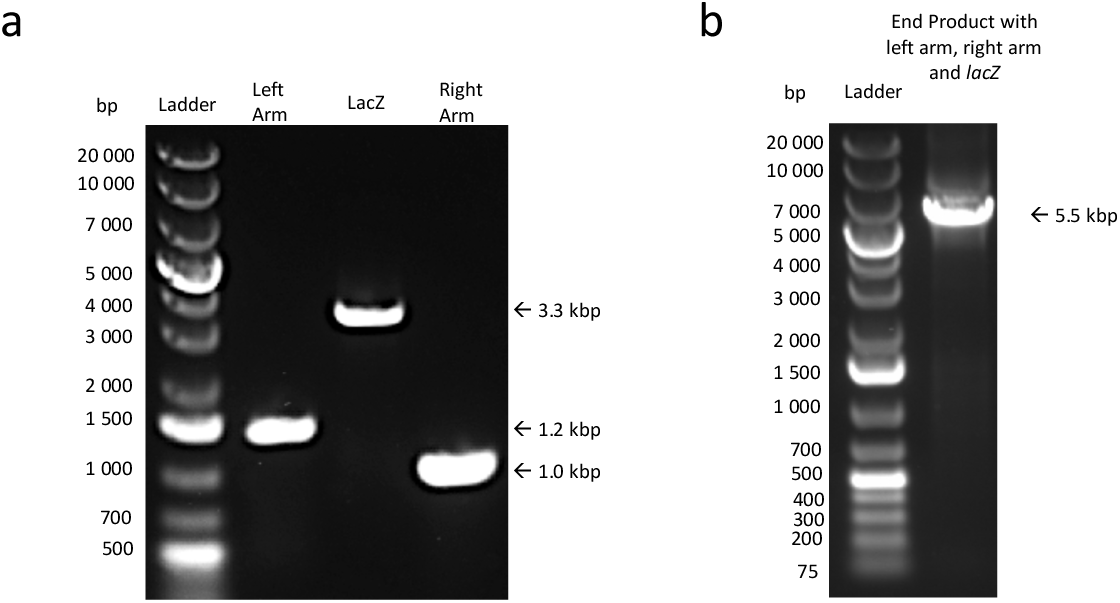
Generation of LacZ construct for *hirL* gene insertion during transformation. Gel Images representing results of PCRs. a) results of extension PCR, where left arm (1.2 kbp), *lacZ* fragment (3.3 kbp) and right arm (1.0 kbp) were obtained. b). Results of SOE-ING PCR, where all three fragments were merged and amplified (final product 5.5 kbp). Arrows indicate size of the obtain products.

**Figure 4.**
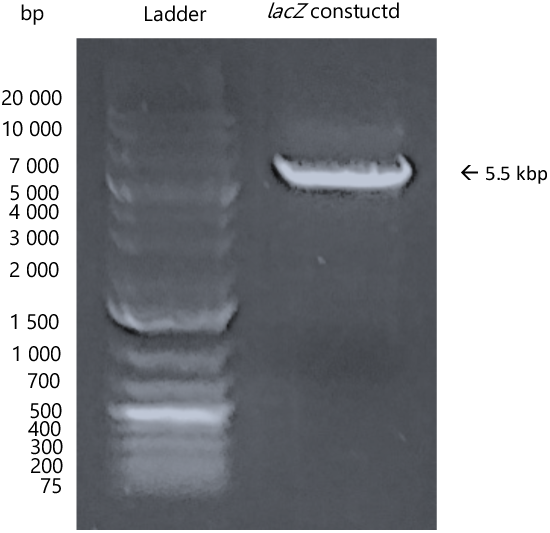
Gel representing fragment of DNA from CP2601 Strain, containing LacZ construct, indicated with arrow (5.5 kbp).

Next, we sought a strain that become competent but does not generate T4P. We removed part of the ComG operon, which encodes proteins required for pilus generation. We have focused on *comGA* and *comGB* since *comGA* encodes a putative ATPase necessary for pilus assembly and encodes a membrane protein. We assumed that partial deletion of those genes would block pilus generation (Figure 5). Based on the sequencing and transformation results, we selected optimal clone for our experiment (Step-by-step strain preparation in experimental procedures). We designed our strain S1672 (Figures 6-7 show the results of the S1672 strain preparation)

**Figure 5.**
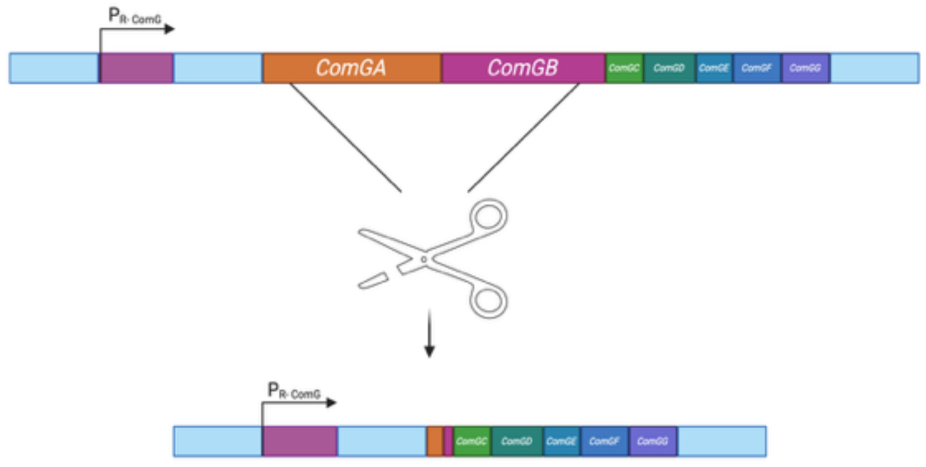
Schematic representation of deletion of *comGA and comGB* genes. LEGEND: P_R-ComG_ - promotor of ComG operon, *ComGA* – *comGA* gene, *ComGB* – *comGB* gene; ComGC – ComGG –rest of ComG operon genes.

**Figure 6.**
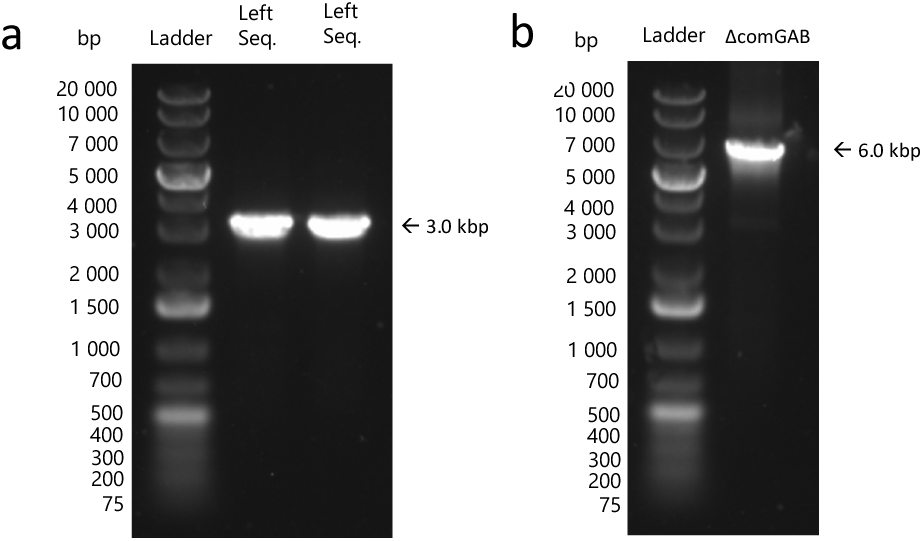
Fragment preparation for deletion of comGAB (ΔcomGAB). a) results of extension PCR, where left arm (3.0 kbp) and right arm (3.0 kbp) were obtained. b). Results of SOE-ING PCR, where both fragments were merged and amplified (final product 6.0 kbp, ΔcomGAB). Arrows indicate size of the obtain products.

**Figure 7.**
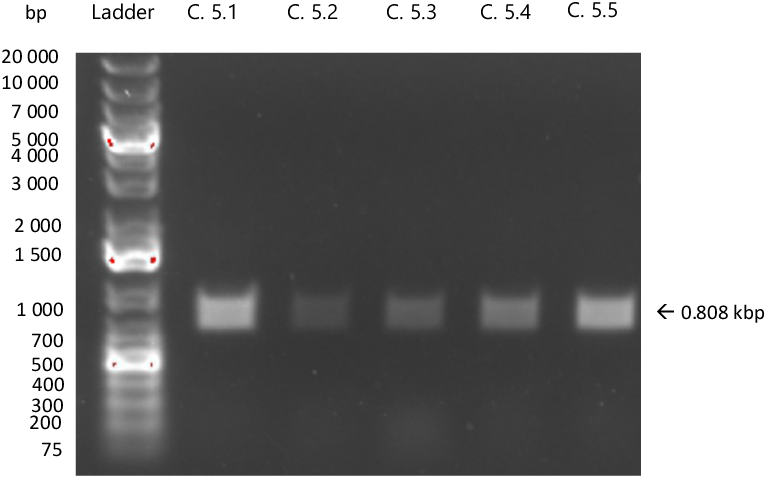
Results of colony PCRs to introduce deletion of *comGAB*. Lines C.5.1 – C.5.5. represent different clones obtained after selection no. 2. Primers used for amplification AB13 and AB14.

### 2. Competence and gene transfer depend on the cell density and medium composition

In parallel with conducting our cell lysis experiments, we identified two additional processes that warranted investigation. One fundamental process is bacterial competence, also known as a quorum sensing mechanism (20). *S. pneumoniae* is known to naturally develop competence and take up extracellular DNA. As previously described, this process involves the sequential activation of early and late competence genes. A key product of the early genes is the competence stimulating peptide (CSP), which initiates a signaling cascade culminating in the expression of late competence genes, including those responsible for pilus formation and DNA uptake. Our competent strains harbor a mutation in the *comA* gene, allowing us to induce competence through exogenous CSP addition, bypassing the defective production of CSP.

In the wild type, competence development in *S. pneumoniae* is cell density dependent. Researchers observed that the logarithmic growth phase, specifically with an optical density (OD) between 0.1 - 0.4, was optimal for competence induction (35), while most of the population becomes refractory to CSP at higher densities (41). However, our observations comport with recent reports by Lam et al. (42), that *S. pneumoniae* can achieve a competent state at more elevated cell densities, if it is harvested and resuspended at a high density in a fresh medium, entering a competent state even at OD 1 (42).

Another critical factor influencing our experiments is the composition of the growth medium. We used chemically defined medium (CDM; see Experimental Procedures for details), supplemented with 5% Todd-Hewitt Broth with Yeast extract (THY). Preliminary testing with various supplements, including CAT and THY media, indicated that 5% THY supplementation yielded the most consistent results (data not shown). Additionally, based on previous reports, we excluded choline chloride from our experimental resuspension medium, since 1% choline chloride inhibits the binding of secreted autolysins such as LytA, LytC and CbpD itself to teichoic and lipoteichoic acids in the pneumococcal cell wall (43) thereby preventing autolysin-mediated cell wall degradation and lysis.

To assess DNA uptake as a measure of competence, we introduced novobiocin-resistant DNA (Nov^R^ DNA) at a final concentration of 0.16 µg/ml following the mixing of attacker and target cells, either in the presence or absence of CSP. After growth in 1× CDM supplemented with 5% THY (containing choline chloride, required for cell growth) to an OD of 0.3, cells were centrifuged and washed three times in the cold 1× CDM + 5% THY (lacking choline chloride), and then resuspended in the same choline-free medium. Cells were incubated for 25 minutes at 37 °C, diluted to 10^-2^, incubated for a 1h at 37 °C, and subsequently plated on selective agar. To quantify transformation events, we plated Nov^R^ CP2204 (Rif^R^) transformants on Rifampicin + Novobiocin (Rif^R^Nov^R^ or RN), while Nov^R^ S1672 (Cm^R^) transformants were selected on Chloramphenicol + Novobiocin (Cm^R^Nov^R^ or CN). The number of colonies growing on these plates represented attacker cells that had successfully acquired Nov^R^ DNA. To calculate transformation efficiency, the number of transformants per ml (CFU/ml, Rif^R^Nov^R^ or Cm^R^Nov^R^) was divided by the total number of attacker cells (CFU/ml, Rif^R^ or Cm^R^, respectively) and multiplied by 100. Table S1 shows number of transformants and Table S2 shows transformation efficiency (CFU/ml or %, respectively, Rif^R^Nov^R^ or Cm^R^Nov^R^) formed during Nov^R^ DNA take up by pilus^+^ CP2204 or pilus^-^ S1672 attackers during competence induction (+CSP) or not (-CSP).

In addition to studying competence development, we explored a second mechanism of horizontal gene transfer, called here gene transfer. Since the T4P pilus is essential for DNA uptake during transformation, its removal from the competent strain prevents this process. Therefore, we specifically examined DNA uptake from lysed cells by competence-induced cells (gene transfer). Håvarstein and colleagues described this process in 2003 (36). They demonstrated that releasing chromosomal DNA into the environment from non-competent cells leads to uptake of the released DNA by competent cells and its integration into their genome via homologous recombination. This process is as important as DNA uptake from environment, since it shows the ability of competent cells to kill and uptake DNA from the neighboring cells, and we are exploring if cell contact is required during that process.

The same strains used in the competence experiments were employed here: attackers-CP2204 (pilus^+^, Rif^R^), S1672 (pilus^-^, Cm^R^), and victim CP2601 (Spc^R^). The gene transfer protocol was identical to that used in the competence assays, with the exception that recovered cells were plated on Rif + Spc (for detecting CP2204 transformants - Rif^R^Spc^R^ or RS) or Cm and Spc (for detecting S1672 transformants - Cm^R^Spc^R^ or CS) to select for cells that had acquired DNA from CP2601. Gene transfer efficiency was calculated using the same formula as transformation efficiency, by dividing the number of transformants per ml (CFU/ml, Rif^R^Spc^R^ or Cm^R^Spc^R^) by the total number of attacker cells (CFU/ml, Rif^R^ or Cm^R^, respectively). Table S3 shows number of transformants and Table S4 shows gene transfer efficiency (CFU/ml or %, respectively, Rif^R^Spc^R^ or Cm^R^Spc^R^) formed during exogenous DNA take up by pilus^+^ CP2204 or pilus^-^ S1672 attackers during competence induction (+CSP) or not (-CSP).

In competence-induced conditions, the pilus^+^ strain CP2204 exhibited decreasing transformation efficiency as OD increased, with the highest transformation efficiency for OD = 0.35 (35%) and lowest for OD = 4 (0.723%) (Figure 8). Our data shows that DNA take up is density-depended but also that the cells can become competent in conditions adjusted for the purposed of cell lysis detection.

**Figure 8.**
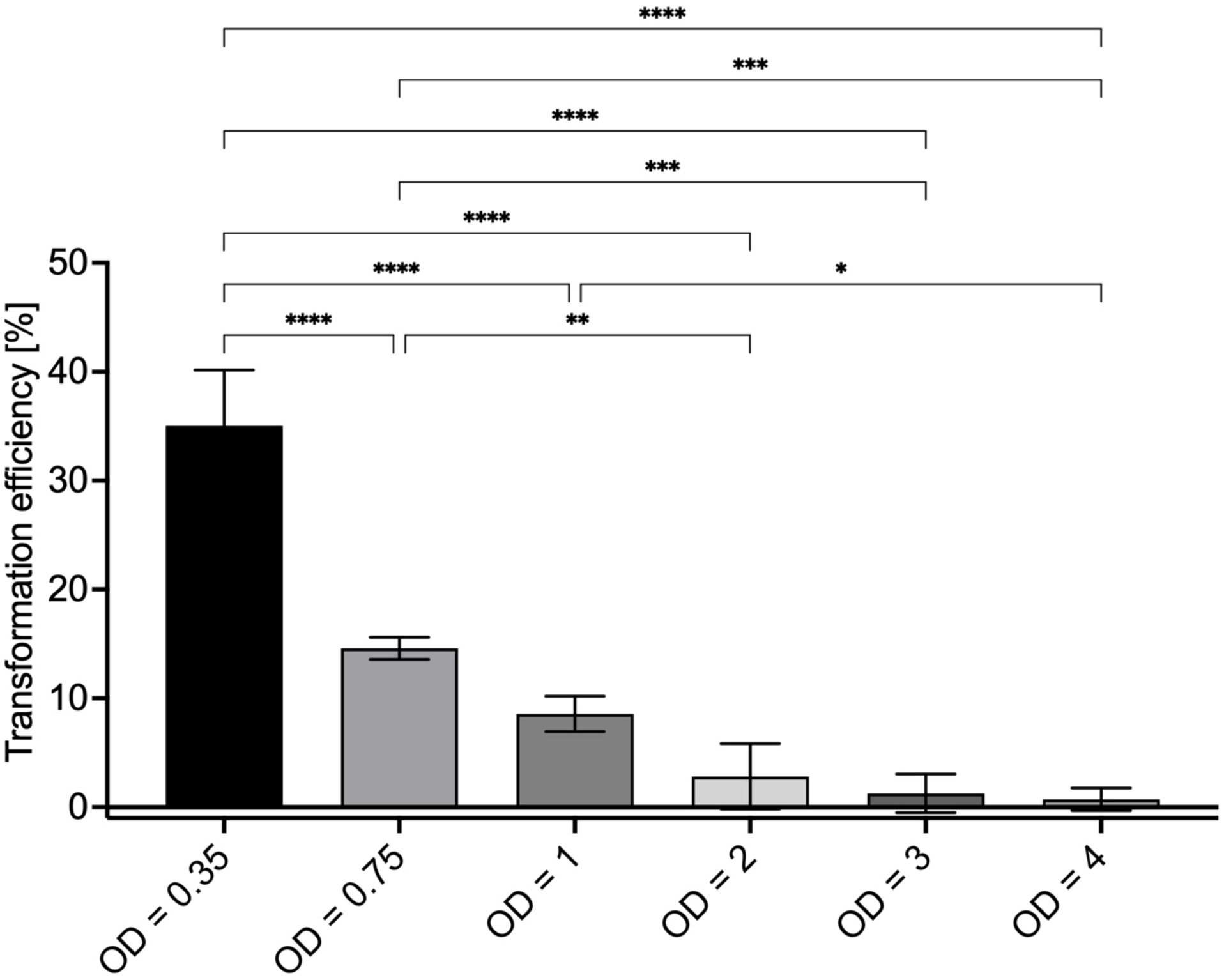
Transformation efficiency of Nov^R^ DNA by pilus^+^ *S. pneumoniae* at different cell densities by pilus^+^ *S. pneumoniae* strain (CP2204 – Rif^R^Nov^R^) under competence-induced conditions (+CSP). The data expresses the mean with corresponding standard deviations, based on results from three independent experiments. Statistical analysis: One-way ANOVA with Tukey’s multiple comparison test, where *p<0.05; **p<0.005, ***p<0.0005, ****p<0.0001

Gene transfer for competence-induced conditions, the pilus^+^ strain CP2204 showed different patter where the highest could be observed for OD = 1 (0.43%) with decreasing for higher or lower ODs, showing that cell density has influence on gene transfer too (Figure 9). Our data shows that for transformation lower ODs is better due to higher DNA accessibility from the environment and for the gene transfer optical OD equal 1 means that cells need higher cells density for cell contact and killing neighbors, but too high cell density can obstruct this process. Again, our data suggest that the cells can become competent, cause lysis of the neighboring cells and take up their DNA in conditions adjusted for the purposed of cell lysis detection.

**Figure 9.**
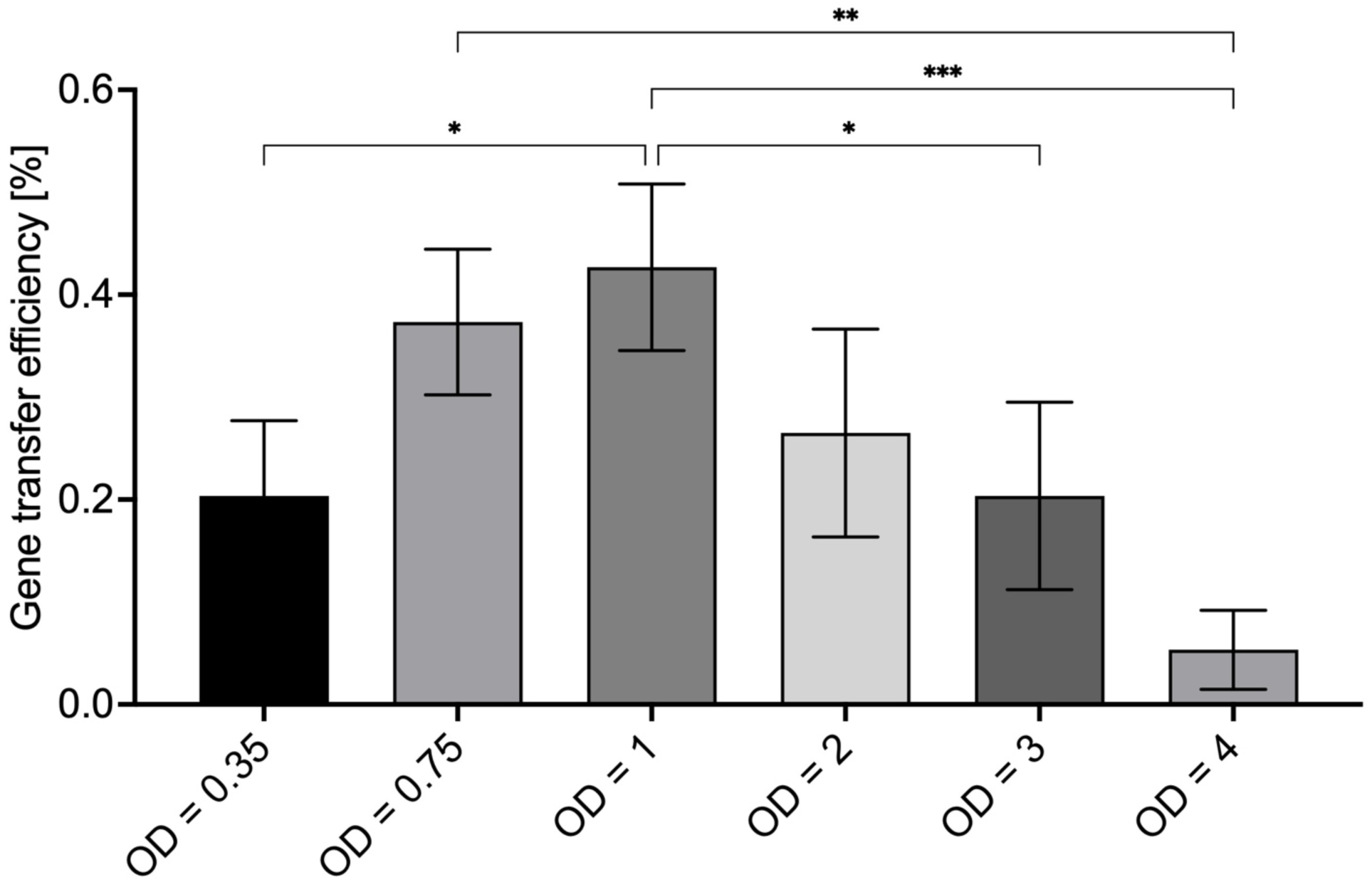
Gene transfer efficiency at different optical cell densities to pilus^+^ *S. pneumoniae* strain (CP2204 – Rif^R^Nov^R^) under competence-induced conditions (+CSP). The data expresses the mean with corresponding standard deviations, based on results from three independent experiments. Statistical analysis: One-way ANOVA with Tukey’s multiple comparison test, where *p<0.05; **p<0.005, ***p<0.0005, ****p<0.0001.

### 3. Non-competent cells release β-galactosidase after contact with CSP-treated cells possessing or lacking *com* T4P

To investigate whether the competence-associated T4P contributes to cell lysis in fratricide, we employed a previously established assay in which the release of β-gal from non-competent cells indicates their lysis induced by competence-induced cells (22,23).

For this assay, we utilized three previously described strains: CP2601 (victim, β-gal^+^), CP2204 (attacker, pilus^+^), and S1672 (attacker, pilus^-^). As mentioned before, β-gal was chosen as a reporter because it remains confined to the cytoplasm and is only released upon cell membrane disruption, thereby serving as a reliable marker of lysis.

Cells were grown in 1x CDM supplemented with 5% THY to an OD of 0.3, incubated on ice for 15–30 minutes, harvested, and washed three times with 1× CDM + 5% THY lacking choline chloride. Cells were resuspended to the specific ODs in the same choline-free medium, as 1% choline chloride inhibits by binding to secreted lysins such as LytA and LytC and consequently, we tested a broader range of cell densities while ensuring that cultures remained in the logarithmic growth phase, during which competence induction is still possible.

Freshly resuspended cells, non-competent and competence-inducible, were mixed at a 1:1 ratio. We combined the cell mixture with an inducer cocktail containing CSP (for competence induction) or left it untreated. Cells were incubated at 37 °C for 25 minutes.

To quantify total β-gal activity, we treated separate samples with 0.1% Triton X-100 and incubated them for 10 minutes at 37 °C to ensure complete lysis. Experimental samples (+/-CSP), were centrifuged and then all the supernatants (+/-CSP, TX100), were transferred to a 96-well plate preloaded with Z-buffer (see Experimental Procedures) which provides the optimal conditions for β-gal enzymatic activity. Following the addition of FDG, we transferred the plate to a plate reader. Fluorescence was recorded every 10 minutes for 90 minutes at 30 °C.

We initially attempted to replicate the 2002 protocol (35), where cells were not harvested and resuspended in a fresh media, just used at OD of range 0.1 – 0.7, but our efforts were unsuccessful. To test whether cell density influences lysis efficiency, we systematically compared 0.35, 0.75, 1, 2, 3, and 4 ODs, since we found out at the preliminary trials that ODs of 1 and 2 failed to produce detectable lysis (data not shown). Contrary to our preliminary experiments, we observed a relationship between cell density and β-gal release, where the highest β-gal release was observed for OD = 1.

To determine if T4P is required for cell lysis we compared pilus^+^ CP2204 and pilus^-^ S1672 strains under competence-induced conditions and observed that there is no statistical significance in β-gal release between fixed OD (for example OD = 1 for both strains) between pilus^+^ and pilus^-^ attackers (Figure 10). Table 1 shows β-gal activity between CSP-treated and not treated samples.

**Figure 10.**
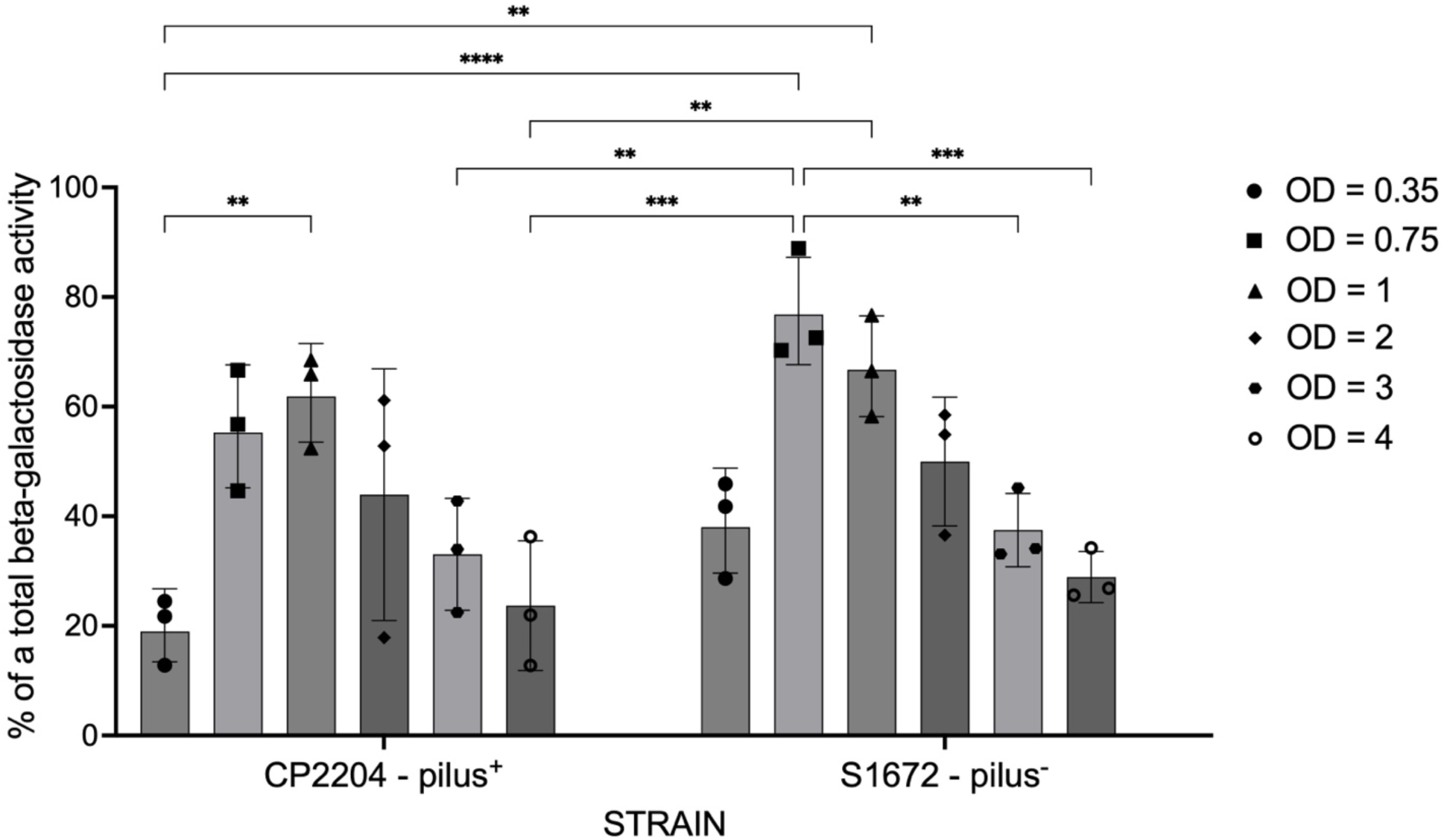
β-Galactosidase Activity released from a victim strain (CP2601) during cell lysis induced by competent *S. pneumoniae* strains at different optical cell densities during cell lysis caused by pilus^+^ attacker (CP2204) or pilus^-^ attacker (S1672), under competence-inducing conditions (+CSP). The data expresses the mean with corresponding standard deviations, based on results from three independent experiments. Statistical analysis: Two-way ANOVA with Tukey’s multiple comparison test, where *p<0.05; **p<0.005, ***p<0.0005, ****p<0.0001.

**Table 1:** β-Gal Activity released from a victim (CP2601) during cell lysis caused by pilus^+^ (CP2204) or pilus^-^ (S1672) *S. pneumoniae* strains under competence-induced conditions (+CSP) at different optical cell densities. The data expresses the mean with corresponding standard deviations, based on results from three independent experiments.

| β-Galactosidase Activity (%) | | | | |
| --- | --- | --- | --- | --- |
| Victim | Attacker | OD | + CSP | - CSP |
| CP2601 – Spc<sup>R</sup><br>hirL::LacZ | CP2204 – pilus<sup>+</sup> | 0.35 | 19.7 ± 6.1 | 1.4 ± 0.7 |
| | | 0.75 | 56.0 ± 11.0 | 1.1 ± 0.4 |
| | | 1 | 62.3 ± 8.6 | 1.2 ± 0.2 |
| | | 2 | 44.0 ± 22.0 | 3.2 ± 0.8 |
| | | 3 | 33.1 ± 10.2 | 3.5 ± 0.9 |
| | | 4 | 23.7 ± 11.9 | 3.8 ± 1.0 |
| | S1672 – pilus<sup>-</sup> | 0.35 | 38.8 ± 9.0 | 1.3 ± 0.9 |
| | | 0.75 | 77.2 ± 10.1 | 1.2 ± 0.7 |
| | | 1 | 67.2 ± 9.2 | 0.9 ± 0.2 |
| | | 2 | 50.0 ± 11.8 | 2.7 ± 1.2 |
| | | 3 | 37.5 ± 6.1 | 4.1 ± 1.6 |
| | | 4 | 28.9 ± 4.7 | 4.1 ± 1.8 |

These results show that the competence T4P is not required for cell lysis under the tested conditions during fratricide. Our findings indicate that an OD of 1 and 0.75 represents the optimal condition for assessing the role of the T4P in competence-induced cell lysis, in the contrast with previous reports of cell lysis disappearing at higher cell densities (35) may reflect their use of grown cultures not cells resuspended in fresh medium.

## EXPERIMENTAL PROCEDURES

### PRIMERS DESIGN

Primers were designed for extension (E-PCR) and splicing by overlapping PCR (SOE-PCR). Table 3 represents primers used to introduce the mutations. Primers were ordered in lyophilized form from Integrated DNA Technologies (IDT) and resuspended to 100 µM in dH_2_O.

### BACTERIAL GROWTH AND MEDIA

Table 3 lists all bacterial strains used in this study. To prepare the required cell stocks, stock of cells with an OD₅₅₀ of 0.2 – 0.3 were cultured in Todd-Hewitt Broth (DIFCO^TM^ Todd-Hewitt Broth, BD) supplemented with yeast extract (BACTO^TM^ yeast extract, Gibco) (THY) until they reached an OD₅₅₀ of 0.2 – 0.3. Adding glycerol (Fisher Scientific) to a final concentration of 15% and aliquoting the cell suspensions into 1 ml cryovials preserved the cells during freezing. Cryovials were frozen using dry ice (–78.5 °C) or placed directly into a –80 °C freezer.

For β-gal assay, cells were grown in CDM (44,45), supplemented with 5% THY until reaching an OD₅₅₀ of 0.3 – 0.4. To run β-gal assays, each strain was washed separately and resuspended in 1x CDM supplemented with 5% THY, with removed choline chloride.

### STRAIN CONSTRUCTION

#### CP2601 construction (β-gal^+^, Spc^R^)

To generate the CP2601 (β-gal^+^, Spc^R^) strain, we employed Extension PCR (E-PCR) and splicing by overlapping PCR (SOE-ing PCR) techniques. We designed primers targeting both flanking regions to create sequences overlapping the *hirL* locus (Table 2). Specifically, primers ABAM1 and ABAM3 were used for the left arm, and primers ABAM4 and ABAM6 for the right arm. We engineered these primers’ 3ʹ and 5ʹ ends to include fragments of the *lacZ* gene, enabling subsequent fusion with the *lacZ* sequence during SOEing-PCR. Following primer design, we performed extension PCR to amplify the overlapping arms and the *lacZ* gene (see below for details). We successfully obtained all required fragments, as shown in Figure 3a.

**TABLE 2:**
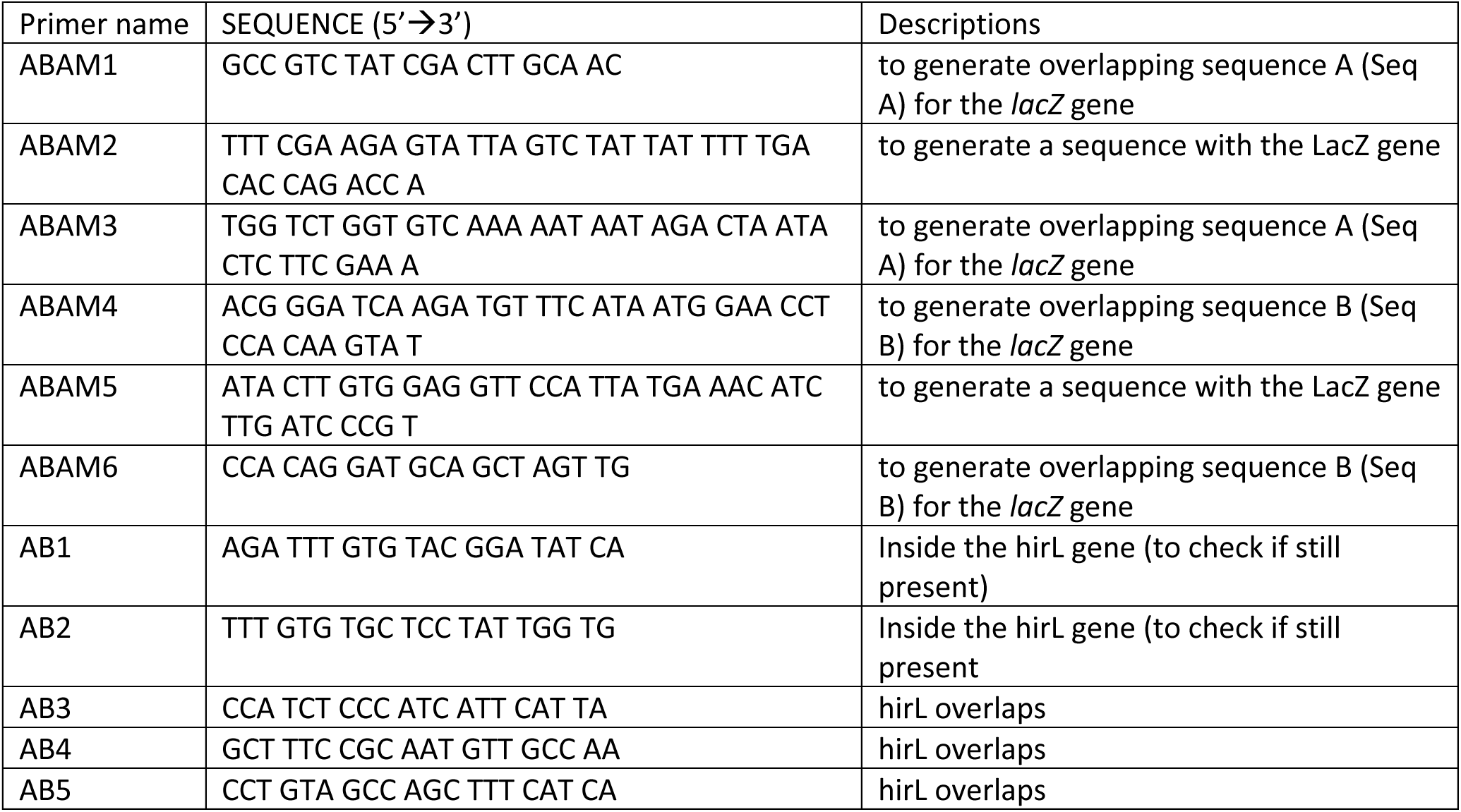
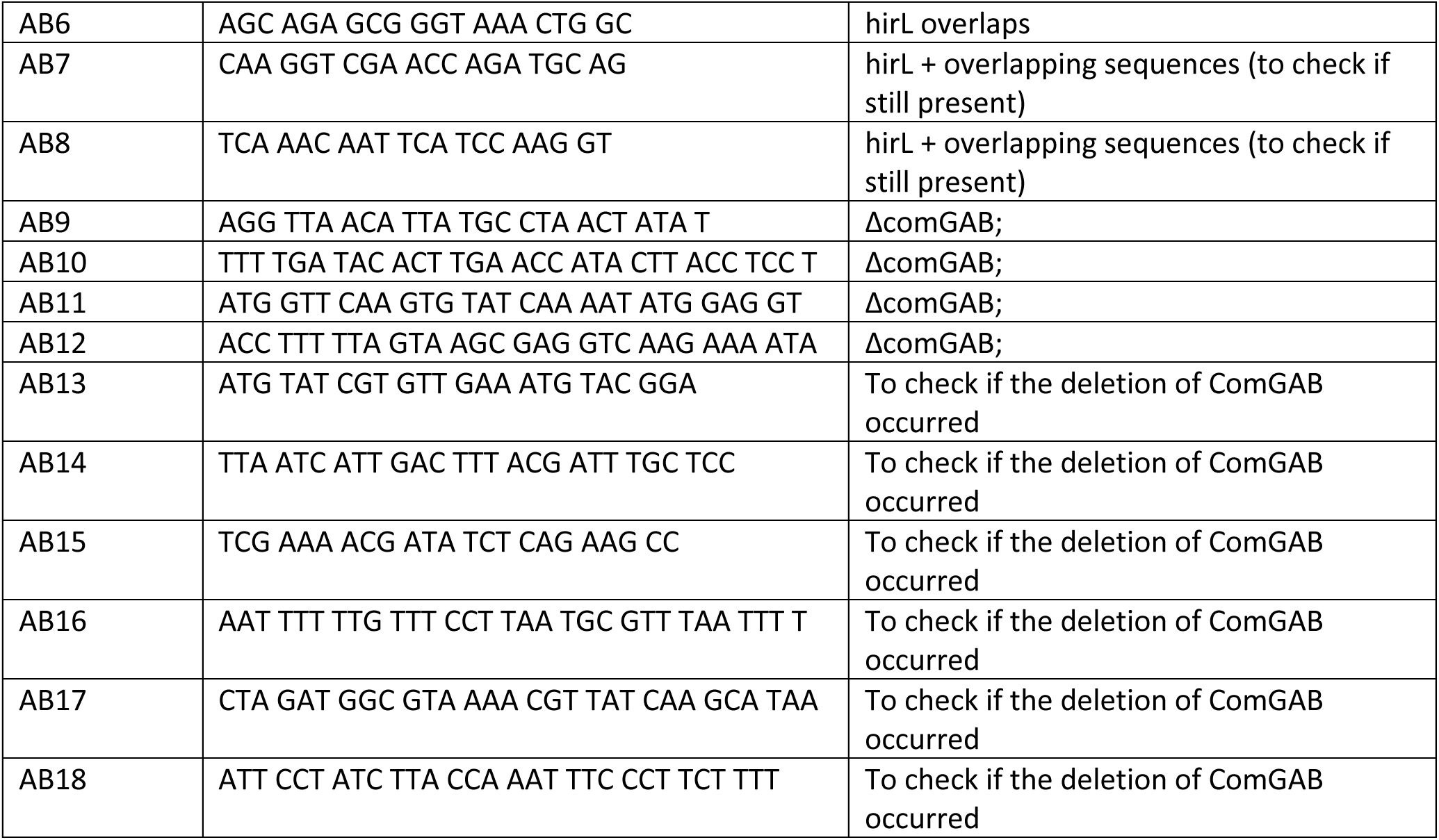
Primers used to introduce mutations to obtain desired strains.

**Table 3:** Bacterial strains used in this study.

| Strain | Description | Reference |
| --- | --- | --- |
| Rx | R36A derivative | (68) |
| CP2000 | Rx but, Δcps cps3D hex cps3D malM511 str-1 bgl-1; Hex<sup>-</sup> Mal<sup>-</sup> Cps<sup>-</sup> Sm<sup>R</sup> Bgal<sup>-</sup> | (69) |
| CP2137 | hex malM511 str-1 bgl-1 ΔcomA cps ssbB[pEVP3]ssbB<sup>+</sup>; Hex<sup>-</sup> Mal<sup>-</sup> ComA<sup>-</sup> Cps<sup>-</sup> SsbB<sup>+</sup> Sm<sup>R</sup> Cm<sup>R</sup> low $\alpha$ -galactosidase background | (70) |
| CP2204 | CP2000 but rpsL1 hlpA::GFP::CAT comA::ermB rif; Hex<sup>-</sup> Mal<sup>-</sup> Sm<sup>R</sup> Cm<sup>R</sup> Cps<sup>-</sup> GFP Em<sup>R</sup> Rif<sup>R</sup> ComA<sup>-</sup> | (11) |
| CP2215 | MD5307 but Δcps, str-1 hlpA::RFP::CAT comE::spc nov-1; Cps<sup>-</sup> Sm<sup>R</sup> Cm<sup>R</sup> Spc<sup>R</sup> Nov<sup>R</sup> Com<sup>-</sup>. | (11) |
| CP2600 | CP2000 but hirL::lacZ; β-Gal<sup>+</sup> | This study. |
| MD5307 | R6 but >10,000 SNPs | (11) |
| CP2601 | CP2601 but comE::Spc<sup>R</sup>; β-Gal<sup>+</sup> Spc<sup>R</sup> | This study. |
| S1671 | CP2137 but comGC::S66C; Hex<sup>-</sup> Mal<sup>-</sup> ComA<sup>-</sup> Cps<sup>-</sup> SsbB<sup>+</sup> Sm<sup>R</sup> Cm<sup>R</sup> low α-galactosidase background | (46) |
| S1672 | S1671 but ΔcomGAB; | This study. |

Following fragment preparation, we performed SOEing-PCR to merge and amplify the three components—the left arm, *lacZ* fragment, and right arm. The reaction successfully generated the desired chimeric fragment of approximately 5.5 kb (Figure 3b).

This recombinant fragment was then introduced into competent *S. pneumoniae* CP2000 cells by transformation (during strain construction, see below). Subsequently, cells were plated on agar (DIFCO^TM^ agar, BD) supplemented with 1.5 % 5-bromo-4-chloro-3-indolyl-β-D-galactopyranoside (X-gal) (ThermoScientific) and incubated for 16 – 24 hours.

Transformation yielded 6.33 × 10^3^ β-gal^+^ transformants, corresponding to a transformation efficiency of 0.18%. Individual blue colonies were picked and inoculated into 4 ml of THY medium. Cultures were incubated overnight (o/n) at room temperature (RT). The following day, cells were transferred to 37 °C and grown until reaching an OD₅₅₀ of 0.2 – 0.5. The three best-performing colonies were selected, frozen, and stored at –80 °C based on growth patterns. The selected colonies were regrown, serially diluted, and plated on X-gal-containing agar plates to obtain a clean clone. After 16 – 24 hours of incubation at 37 °C, we picked ten individual colonies from each of the three selected candidates, resuspended in 4 ml of THY, and incubated o/n at RT, followed by growth at 37 °C the next day. Based on growth characteristics and the consistent presence of only blue colonies upon re-plating, we selected colony 2.2 for DNA purification and sequencing, designated here CP2600.

To generate the final non-competent strain, we first purified genomic DNA from strain CP2215 using an ethanol-based DNA extraction protocol (see details below). This DNA served as the template for PCR amplification of the *comE::Spc* region. We used the PCR product to transform the previously engineered strain harboring the *lacZ* gene.

Briefly, CP2600 were grown to an OD_550_ of 0.05 – 0.1, mixed with CSP, BSA, and CaCl₂, and incubated for 1 hour at 37°C to promote homologous recombination. Following incubation, cells were plated on agar containing X-gal and spectinomycin (Spc). This transformation yielded 3.46 × 10^5^ β-gal^+^, Spc-resistant (Spcᴿ) transformants, corresponding to a transformation efficiency of 0.99%.

The same selection was performed on Spc^R^ mutants as for *lacZ* as described above. To verify the resulting strain’s loss of competence, a control transformation was conducted using Novobiocin-resistant (Nov^R^) DNA. No transformants were detected, with a transformation efficiency below 0.00001%, confirming the strain’s non-competent phenotype. The final β-gal^+^, Spcᴿ, non-competent strain was designated CP2601.

#### S1672 Construction

As with the generation of strain CP2601, we designed primers to overlap with the *comGA* and *comGB* regions while avoiding amplifying the full *comGA–comGB* sequence. We used Primers AB9 and AB10 to amplify the left homology arm, which included a sequence overlapping comGA and a short region complementary to the residual comGB sequence. Similarly, primers AB11 and AB12 were employed to amplify the right homology arm, designed to include overlap with the remaining comGA sequence and complementarity to the region adjacent to comGB. We performed Extension PCR to generate the left and right arms (Figure 6a). To enable deletion of the target region, the two arms were subsequently fused using overlapping sequences through splicing by overlap extension PCR (Figure 6b). To generate the ΔcomGAB mutant, a deletion construct was introduced into the competent S1671 strain, which Lam et al previously used to visualize T4P (46) The transformation protocol followed the same approach used for generating the β-gal^+^ strain. Successful integration of the deletion was verified by PCR using primers AB13 and AB14. In the wild type, this primer pair amplified a 2,995 bp fragment. In contrast, in the ΔcomGAB strain, a shortened 808 bp product was detected, confirming the successful removal of the targeted comGA–comGB region.

Following transformation, we picked single colonies, performed colony PCR, and streaked them on an agar plate. Plates were incubated for 16 – 24h at 37°C, and the results of colony PCR were observed on 0.8% agarose (GoldBio) gel. Gel analysis revealed a mixed population, with some cells harboring the targeted comGAB deletion (808 bp fragment) and others retaining the wild-type allele (3 kbp fragment), as observed in colony 14 (Figure 6).

The three previously selected colonies (14,19,21) were each inoculated into 4 ml of THY, serially diluted (from 10⁰ to 10⁻³), and incubated o/n at RT, followed by growth at 37 °C, until they have reached OD_550_ = 0.3-0.5 . We subsequently froze cultures for future selection steps. Colonies 14, 19, and 21—chosen based on optimal growth—were cultivated at 37 °C, serially diluted, and plated (all from the 10⁻¹ dilution due to superior growth performance). After 16 – 24 hours of incubation at 37 °C, we picked ten colonies from each plate, resuspended them in 4 ml of THY, and prepared 50 µl colony PCR mixtures. Three promising clones were selected based on growth in liquid culture and PCR results. These clones were grown again, serially diluted, and plated. We picked five single colonies from each of the three chosen clones, resuspended them in 4 ml of THY, and subjected them to colony PCR (Figure 7). This process identified three colonies as potential candidates for the desired strain (strong band on the agarose gel). Genomic DNA from these three clones was purified and submitted for sequencing to confirm the intended genetic modification. Based on sequencing confirmation and growth characteristics, we selected clone C.5.5 for subsequent experiments and named S1672.

### EXTENSION POLYMERASE CHAIN REACTION (E-PCR)

Extension PCR was performed to generate fragments for overlapping or deletion PCR. Phire^TM^ Reaction Buffer Kit (ThermoScientific) and dNTP Mix (ThermoScientific) were used. For *lacZ* and its overlapping regions (Seq A and Seq B), primers ABAM1–ABAM6 (listed in Table 2) were used to introduce overlapping sequences at the fragment ends. The PCR protocol consisted of an initial denaturation step (3 min at 98 °C), followed by 30 cycles of denaturation (8 s at 98 °C), annealing (8 s at 64 °C for Seq A and Seq B, and 65 °C for LacZ), and extension (12 s for Seq A and Seq B, and 33 s for LacZ at 72 °C), concluding with a final extension step (3 min at 72 °C).

Two overlapping fragments were generated for deletion PCR with matching ends that excluded the targeted sequence. The procedure was similar, except primers AB9–AB12 were used, we adjusted the annealing temperature to 55 °C, and the extension times to 30 and 31 seconds.

We analyzed all PCR products on a 0.8% agarose gel and purified them using a commercially available PCR clean-up kit (Zymo Research, DNA Clean & Concentrator^TM^-25 ).

### SPLICING BY OVERLAPPING POLYMERASE CHAIN REACTION (SOE-PCR)

To assemble the prepared Seq A, LacZ, and Seq B fragments, Splicing by Overlap Extension PCR (SOE-ing PCR) was employed. Phire^TM^ Reaction Buffer Kit (ThermoScientific) and dNTP Mix (ThermoScientific) were used. In the first step, we carried out overlap extension PCR (OL-PCR) without primers. The protocol included an initial denaturation (3 min at 98 °C), followed by 15 cycles of denaturation (8 s at 98 °C), annealing (8 s at 60 °C), and extension (55 s at 72 °C), ending with a final extension step (3 min at 72 °C).

In the second step of OL-PCR, we amplified the newly formed fragment using primers ABAM1 and ABAM6. This step followed a similar protocol: initial denaturation (3 min at 98 °C), 20 cycles of denaturation (8 s at 98 °C), annealing (8 s at 64 °C), and extension (55 s at 72 °C), followed by a final extension (3 min at 72 °C).

We utilized SOE-ing PCR to delete portions of the *comG* operon, which encodes pilus proteins. Primers AB9–AB12 (listed in Table 2) were used to remove segments of comGA and comGB. The first step (extension PCR) followed the protocol described previously. The second step included initial denaturation (3 min at 98 °C), 30 cycles of denaturation (8 s at 98 °C), annealing (8 s at 60 °C), and extension (62 s at 72 °C), concluding with a final extension (3 min at 72 °C).

We analyzed all PCR products on a 0.8% agarose gel, purified using a commercially available PCR clean-up kit (Zymo Research, DNA Clean & Concentrator^TM^-25), and subsequently introduced into *S. pneumoniae* strains via transformation, as detailed below.

### COLONY POLYMERASE CHAIN REACTION (PCR)

Colony PCR was performed to confirm the presence of newly introduced sequences after transformation. Phire^TM^ Reaction Buffer Kit (ThermoScientific) and dNTP Mix (ThermoScientific) were used. A single colony was picked from the plate with an inoculation needle and transferred to 4 ml of THY (for growth and stock). After that the same needle with the same colony was moved into a PCR tube containing 50 µl of PCR mix (10 µl of 5x reaction buffer, 1 µl of 10 mM dNTPs mix, 2.5 µl of 10 µM primer 1, 2.5 µl of 10 µM primer 2, 1 µl of Phire II polymerase, water). Additionally, the colony was streaked onto an agar plate to allow further growth in preparation for subsequent colony expansion.

Protocol included: initial denaturation (3 min @ 98°C), denaturation (8 s @ 98°C), annealing (8 s, 60°C), extension (30 s @ 72°C), where denaturation, annealing, and extension were repeated in 20 cycles and final extension (3 min @ 72°C). We analyzed all PCR products on a 0.8% agarose gel.

### DNA PURIFICATION

Ethanol precipitation was performed to purify genomic DNA from CP2000 (*hirL*) or CP2215 (comE:;spc), which served as the template for mutagenesis. Cells were cultured in 12 ml of THY medium until reaching an OD_550_ of 0.4. EDTA (Quality Biological) was added to a final concentration of 0.5 mM, followed by incubation on ice for 30 minutes. The cells were then centrifuged (7000 rpm, 7 min, 4°C) and resuspended in a buffer containing 0.1 M NaCl (VWR), 0.01 M Tris-HCl (Invitrogen), and 5 mM EDTA, supplemented with 0.2% Triton X-100 (MP) and 1% RNase (2 mg/ml). This mixture was incubated at 37°C for 20 minutes. SDS (Fisher BioReagents) was added to a final concentration of 1%, and the solution was incubated at 55°C for an additional 20 minutes. Sodium acetate (Ambion) was added to reach a final concentration of 0.3 M, and the sample was frozen at –80°C overnight or processed immediately.

DNA was extracted by adding an equal volume of extraction buffer (chloroform (Fisher Chemical) mixed with 4% isoamyl alcohol (Fisher Scientific) mixed), and centrifuged (8000 × g for 30 min or 12,000 × g for 10 min at room temperature). The aqueous phase was transferred to a fresh tube, and the extraction was repeated using the same buffer and centrifugation steps. After transferring the upper phase to a new tube, an equal volume of pre-chilled (–20°C) isopropanol (Fisher Chemical) was added, followed by vortexing and centrifugation (7000 × g for 15 min or 12,000 × g for 5 min at 4°C). The pellet was washed with 500 µl of cold wash buffer (75% ethanol (Fisher Bioreagents), 3% sodium acetate, stored at 4°C), vortexed, and centrifuged (12,000 × g, 5 min, 4°C). This wash step was repeated two more times. After drying the pellet, DNA was resuspended in 50 µl of TE buffer (10 mM Tris (Fisher Bioreagents), 1 mM EDTA, pH 8.0), vortexed, and the concentration was measured.

### COMPETENCE INDUCTION

Three components are required to induce competence in *S. pneumoniae*. The first is bovine serum albumin (BSA, GoldBio), prepared as a 4 mg/ml (0.4%) stock solution in distilled water and stored at 4°C. The second is calcium chloride (CaCl₂, Ward’s Science), prepared as a 0.1 mM stock solution in distilled water and kept at room temperature. The third component is CSP (Neo BioLab), stored at –20°C as a 100 µg/ml stock solution in distilled water. We prepared temporary working stocks of 0.4% BSA and 0.01 M CaCl₂ in water for inducer preparation. The final inducer mix consisted of 20 µl of 0.4% BSA, 100 µl of 0.01 M CaCl₂, 40 µl of 100 µg/ml CSP, and 840 µl of culture medium

#### TRANSFORMATION: During strain construction

To introduce specific mutations into bacterial strains, cells were cultured in 10 ml of THY medium (pH 6.6) until reaching an optical density at 550 nm (OD₅₅₀) of 0.05–0.1. Cultures were then diluted 1:1000 for LacZ insertion or *comGAB* deletion, and 1:10 for insertion of the *Spcᴿ* gene. A total of 1 ml of the diluted culture was mixed with 10 µl of 100 ng/ml CSP, 10 µl of 4% BSA, and 10 µl of 0.1 M CaCl₂. DNA (LacZ insert, Δ*comGAB* construct, or genomic DNA from CP2215 carrying *Spcᴿ*) was added to a final 160 ng/ml concentration. Mixtures were incubated at 37 °C for 3 hours (LacZ and Δ*comGAB*) or 1 hour (*Spcᴿ*). After incubation, cultures were serially diluted in THY and plated on selective agar plates: X-gal-containing plates for LacZ screening, X-gal and Spc-containing plates for *Spcᴿ* selection, and non-selective plates for Δ*comGAB* due to the absence of a selectable marker. After 16–24 hours of incubation, blue colonies (LacZ⁺), *Spcᴿ*colonies, or randomly selected colonies (Δ*comGAB*) were picked to confirm the corresponding gene insertion or deletion.

#### TRANSFORMATION: Assessment of competence or gene transfer

To assess whether cells became competent or whether the attacker strain acquired genes from the victim, cultures were grown in 1x CDM + 5% THY to OD OD_550_ of 0.3. Cells were withdrawn, incubated on ice for at least 15 minutes, centrifuged, and washed three times with 5 ml of 1× CDM + 5% THY lacking choline chloride. Cells were resuspended to a specific OD_550_ depending on the target final OD: for OD_550_ values of 0.35, 0.75, 1, 2, 3, 4 cells were adjusted to OD_550_ 1.4, 3, 4, 8 and 16, respectively. Attacker and victim strains were mixed in a 1:1 ratio and combined with an inducer cocktail (containing BSA, CaCl₂, and CSP or no CSP) at a 1:1 ratio. The mixture was incubated at 37°C for 25 minutes, serially diluted (10⁻² to 10⁻⁷), and plated on agar plates containing Rif (Sigma Aldrich) and Nov (Sigma Aldrich) or Cm (GoldBio) and Nov (for competence assessment) or Rif and Spc (Sigma Aldrich) or Cm and Spc (for gene transfer detection). Plates were incubated for 40 – 48 hours at 37 °C.

### BETA-GALACTOSIDASE ASSAY

Cells were cultivated in 1× CDM supplemented with 5% THY until reaching an optical density at 550 nm (OD₅₅₀) of 0.3. Cultures were then harvested, incubated on ice for a minimum of 15 minutes, centrifuged, and washed three times with 5 ml of 1× CDM + 5% THY lacking choline chloride. The cell pellets were resuspended in the same medium to achieve target OD₅₅₀ values of 1.4, 3, 4,8, and 16, corresponding to final OD₅₅₀ values of 0.35, 0.75, 1, 2, 3 and 4, respectively. Cell suspensions were mixed at a 1:1 ratio based on OD, then an equal volume of inducer cocktail containing BSA, CaCl₂, and either CSP or no CSP was added.

A final concentration of 160 ng/µl of Nov^R^ DNA was added to this mixture. The reaction mixtures were incubated at 37°C in a thermoblock for 25 minutes, then rapidly cooled on ice with water. The reaction was split for samples treated with CSP to quantify total enzyme release. Triton X-100 was added for cell lysis, and 0.1% of each reaction was incubated at 37°C for 10 minutes.

Samples (with CSP, without CSP, and lysed controls) were then transferred to a microplate containing Z-buffer (0.3 M Na₂HPO₄ (Sigma Aldrich), 0.2 M NaH₂PO₄ (Sigma), 50 mM KCl (STERM CHEMICALS INC), 5 mM MgSO₄ (ACROS ORGANICS), and 0.25 M β-mercaptoethanol (Sigma Aldrcih)) supplemented with 50 µM FDG (Invitrogen). Fluorescence was measured at 30°C over 90 minutes.

## DISCUSSION

Here, we present the involvement of competence T4P of *S. pneumoniae* in cell lysis induced by fratricide. Our data show that deletion of T4P genes did not cause a decrease in β-gal release from lysed victim cells during fratricide; instead, we observed an increase in β-gal release. This effect may be due to changes in cell surface properties and an enhanced role of lytic proteins such as LytA, LytC, and CbpD, which—following pilus removal—could display increased lytic activity. Considering our results alongside the known role of lysins in cell lysis, we conclude that fratricide-induced lysis depends on the presence of these lytic enzymes rather than the pilus itself. However, researchers still need to confirm the requirement for close contact, as CbpD acts during direct cell interactions. Additionally, we observed a non-linear relationship between OD and β-gal activity, with OD = 1 producing the highest activity, suggesting a parabolic rather than a linear trend. While our findings indicate that T4P is not essential for cell lysis, they confirm previous reports that T4P is crucial for DNA uptake during transformation and for gene transfer events associated with fratricide.

Beyond *S. pneumoniae*, T4P has been described in numerous bacteria, including *Pseudomonas aeruginosa* (47), *Vibrio cholerae* (48,49), *Bacillus subtilis* (*28*), *Acinetobacter baylyi* (50,51), and *Neisseria* spp (52). including *N. gonorrhoeae* (53). In all these species, T4P is involved in DNA binding during transformation. Before the discovery of T4P in *S. pneumoniae* in 2013, the Håvarstein group described fratricide between *S. pneumoniae* and closely related species such as *Streptococcus mitis* and *Streptococcus oralis* (39). Fratricide-induced cell lysis and DNA release have also been reported in *Enterococcus faecalis* (54,55), *Staphylococcus aureus*, *Streptococcus epidermidis* (56), and *Streptococcus mutans* (57,58). Of these, only *S. mutans* involves T4P in fratricide, though the pathways differ between *S. pneumoniae* and *S. mutans* (*57*). Moreover, *S. mutans* can release DNA via membrane vesicles (59). Other bacteria possessing T4P release DNA through mechanisms distinct from fratricide-induced lysis. For example, *P. aeruginosa* (47)—one of the most studied species— employs four DNA release strategies: phenazine-and pyocyanin-induced autopoisoning (60,61), HQNO autopoisoning (62), prophage endolysin activation (63,64), and explosive cell lysis (63). *B. subtilis* (28) produces T4P but releases DNA without cell lysis (65), while *N. gonorrhoeae* relies on a type IV secretion system for DNA release (66,67). Despite the widespread occurrence of T4P across bacterial species, its involvement in fratricide-induced cell lysis has not been reported beyond *S. mutans*.

Extended experimental troubleshooting led us to optimize conditions for studying fratricide-associated cell lysis, ultimately using a medium without choline. During this process, we tested several media, including THY, 1× CDM supplemented with 5% THY, and CDM with CAT. None of these conditions resulted in β-gal release, even when the pilus was present. As noted previously, even as little as 1% choline (or lower concentrations) can inhibit lysin-dependent lysis. Since our findings indicate that fratricide-associated lysis depends on multiple lysins, this raises an important question: how does this process occur *in vivo*, where choline levels are uncontrolled and cells grow within biofilms? Future studies should investigate whether deleting genes encoding minor subunits of T4P produces a different effect. Direct visualization of fratricide, which is another goal of our ongoing work, could provide further insights. Moreover, conducting experiments under biofilm conditions—better reflecting *in vivo* environments compared to *in vitro*—may yield more physiologically relevant data. As for our goal, determining how the cell lysis process during fratricide occurs could, in the future, help identify novel therapeutic targets for vaccines or provide new strategies to prevent the horizontal gene transfer of antibiotic-resistance genes.

