## Supplementary material for "*Streptococcus pneumoniae* fratricide-induced cell lysis does not require the type IV competence pilus": n/a

| **Number of Nov$^R$ Transformatns (CFU/ml)** | | | | |
|---|---|---|---|---|
| DNA | Attacker | OD | + CSP | - CSP |
| Nov$^R$ DNA | CP2204 – pilus$^+$ | 0.35 | 6.3 x $10^7$ ± 2.8 x $10^7$ | 0.6 x $10^1$ ± 1.0 x $10^1$ |
| | | 0.75 | 4.8 x $10^7$ ± 1.2 x $10^7$ | 2.2 x $10^1$ ± 2.6 x $10^1$ |
| | | 1 | 5.5 x $10^7$ ± 3.6 x $10^7$ | 7.4 x $10^1$ ± 9.5 x $10^1$ |
| | | 2 | 1.5 x $10^7$ ± 7.3 x $10^6$ | 1.3 x $10^2$ ± 2.3 x $10^2$ |
| | | 3 | 9.1 x $10^6$ ± 8.4 x $10^6$ | 4.6 x $10^2$ ± 8.0 x $10^2$ |
| | | 4 | 8.6 x $10^6$ ± 9.7 x $10^6$ | 7.2 x $10^2$ ± 1.1 x $10^3$ |
| | S1672 – pilus$^-$ | 0.35 | 9.3 x $10^3$ ± 1.0 x $10^4$ | 1.3 x $10^3$ ± 2.1 x $10^3$ |
| | | 0.75 | 6.1 x $10^4$ ± 4.6 x $10^4$ | 2.2 x $10^2$ ± 2.6 x $10^2$ |
| | | 1 | 9.9 x $10^4$ ± 6.5 x $10^4$ | 2.8 x $10^2$ ± 2.6 x $10^2$ |
| | | 2 | 3.5 x $10^4$ ± 2.9 x $10^5$ | 1.6 x $10^3$ ± 2.0 x $10^3$ |
| | | 3 | 5.6 x $10^4$ ± 6.3 x $10^3$ | 6.3 x $10^3$ ± 4.7 x $10^3$ |
| | | 4 | 3.3 x $10^4$ ± 2.1 x $10^4$ | 1.8 x $10^4$ ± 2.6 x $10^4$ |

**Table S1:** Number of Transformants per ml (CFU/ml) that were formed during Nov$^R$ DNA uptake by pilus$^+$ (CP2204) or pilus$^-$ (S1672) *S. pneumoniae* strains when competence was induced (+CSP) or not (-CSP) at different optical cell densities. The data are expressed as means with corresponding standard deviations, based on results from three independent experiments.

| **Transformation efficiency (%)** | | | | |
|---|---|---|---|---|
| DNA | Attacker | OD | + CSP | - CSP |
| Nov$^R$ DNA | CP2204 – pilus$^+$ | 0.35 | 35.0 ± 5.1 | <0.0001 |
| | | 0.75 | 14.6 ± 1.0 | <0.0001 |
| | | 1 | 8.6 ± 1.6 | <0.0001 |
| | | 2 | 2.828 ± 3.0 | <0.0001 |
| | | 3 | 1.275 ± 1.8 | <0.0001 |
| | | 4 | 0.723 ± 1.0 | <0.0001 |
| | S1672 – pilus$^-$ | 0.35 | <0.1 | <0.0001 |
| | | 0.75 | <0.1 | <0.0001 |
| | | 1 | <0.1 | <0.0001 |
| | | 2 | <0.01 | <0.001 |
| | | 3 | <0.01 | <0.001 |
| | | 4 | <0.01 | <0.001 |

**Table S2:** Transformation efficiency of Nov$^R$ DNA uptake by pilus$^+$ (CP2204) or by pilus$^-$ (S1672) *S. pneumoniae* strains when competence was induced (+CSP) or not (-CSP) at different optical cell densities. The data are expressed as means with corresponding standard deviations, based on results from three independent experiments.

| Number of $Spc^R$ Transformants (CFU/ml) | | | | |
|---|---|---|---|---|
| Deficient strain | Attacker | OD | + CSP | - CSP |
| CP2601 – $Spc^R$ hirL::LacZ | CP2204 – $pilus^+$ | 0.35 | $3.8 \times 10^5 \pm 2.3 \times 10^5$ | $4.1 \times 10^2 \pm 1.3 \times 10^2$ |
| | | 0.75 | $1.3 \times 10^6 \pm 5.0 \times 10^5$ | $4.6 \times 10^2 \pm 1.4 \times 10^2$ |
| | | 1 | $1.8 \times 10^6 \pm 8.3 \times 10^5$ | $5.8 \times 10^2 \pm 1.5 \times 10^2$ |
| | | 2 | $2.1 \times 10^6 \pm 9.9 \times 10^5$ | $1.8 \times 10^3 \pm 1.4 \times 10^3$ |
| | | 3 | $2.3 \times 10^6 \pm 1.0 \times 10^6$ | $1.7 \times 10^3 \pm 1.3 \times 10^3$ |
| | | 4 | $9.3 \times 10^5 \pm 7.3 \times 10^5$ | $1.9 \times 10^3 \pm 1.6 \times 10^3$ |
| | S1672 – $pilus^-$ | 0.35 | $1.1 \times 10^2 \pm 1.9 \times 10^2$ | $8.3 \times 10^2 \pm 1.4 \times 10^2$ |
| | | 0.75 | $2.2 \times 10^3 \pm 2.6 \times 10^3$ | $7.2 \times 10^2 \pm 6.3 \times 10^2$ |
| | | 1 | $2.7 \times 10^3 \pm 7.7 \times 10^2$ | $1.2 \times 10^3 \pm 1.9 \times 10^3$ |
| | | 2 | $8.2 \times 10^3 \pm 4.7 \times 10^3$ | $3.4 \times 10^4 \pm 6.9 \times 10^4$ |
| | | 3 | $4.6 \times 10^3 \pm 8.6 \times 10^2$ | $2.3 \times 10^4 \pm 3.9 \times 10^4$ |
| | | 4 | $6.2 \times 10^3 \pm 5.1 \times 10^3$ | $8.6 \times 10^4 \pm 1.5 \times 10^5$ |

**Table S3:** Number of transformants per ml (CFU/ml) that were formed during $Spc^R$ gene transfer from the victim (CP2601) to $pilus^+$ (CP2204 - $Rif^R$) or $pilus^-$ (S1672 - $Cm^R$) *S. pneumoniae* strains when competence was induced (+CSP) or not (-CSP) at different optical cell densities. The data are expressed as means with corresponding standard deviations, based on results from three independent experiments.

| Gene Transfer efficiency (%) | | | | |
|---|---|---|---|---|
| Deficient strain | Attacker | OD | + CSP | - CSP |
| CP2601 – $Spc^R$ hirL::LacZ | CP2204 – $pilus^+$ | 0.35 | $0.20 \pm 0.07$ | <0.0001 |
| | | 0.75 | $0.37 \pm 0.07$ | <0.0001 |
| | | 1 | $0.43 \pm 0.08$ | <0.0001 |
| | | 2 | $0.256 \pm 0.1$ | <0.001 |
| | | 3 | $0.204 \pm 0.9$ | <0.001 |
| | | 4 | $0.053 \pm 0.4$ | <0.001 |
| | S1672 – $pilus^-$ | 0.35 | <0.00001 | <0.00001 |
| | | 0.75 | <0.00001 | <0.00001 |
| | | 1 | <0.00001 | <0.00001 |
| | | 2 | <0.0001 | <0.0001 |
| | | 3 | <0.0001 | <0.0001 |
| | | 4 | <0.0001 | <0.0001 |

**Table S4:** Gene transfer efficiency of $Spc^R$ gene by $pilus^+$ (CP2204 - $Rif^R$) or $pilus^-$ (S1672 - $Cm^R$) *S. pnuemoniae* strains when competence was induced (+CSP) or not (-CSP) at different optical cell densities. The data are expressed as means with corresponding standard deviations, based on results from three independent experiments.
